# A coordinated switch in sucrose and callose metabolism enables enhanced symplastic unloading in potato tubers

**DOI:** 10.1101/2023.11.24.568555

**Authors:** Bas van den Herik, Sara Bergonzi, Yingji Li, Christian W. Bachem, Kirsten H. ten Tusscher

**Affiliations:** Computational Developmental Biology, Utrecht University, Utrecht, Netherlands; Plant Breeding, Wageningen University & Research, Wageningen, The Netherlands

## Abstract

One of the early changes upon tuber induction is the switch from apoplastic to symplastic unloading. Whether and how this change in unloading mode contributes to sink-strength has remained unclear. In addition, developing tubers also change from energy to storage-based sucrose metabolism. Here we investigated the coordination between changes in unloading mode and sucrose metabolism and their relative role in tuber sink strength by looking into callose and sucrose metabolism gene expression combined with a model of apoplastic and symplastic unloading. Decreased callose deposition in tubers is driven by decreased callose synthase activity. Furthermore, changes in callose metabolism and sucrose metabolism are strongly correlated, indicating a well-coordinated developmental switch. Modelling indicates that symplastic unloading is not the most efficient unloading mode per se. Instead, it is the concurrent metabolic switch that provides the physiological conditions necessary to potentiate symplastic transport and thereby enhance tuber sink strength.

## Introduction

In potato, the tuberigen *StSP6A* induces tuber onset (Navarro et al., 2011) which is a major developmental transition, associated with large changes in plant physiology among which is the emergence of a new, and strong, sucrose sink. Besides its role in tuber establishment by switching on the tuber developmental program, *StSP6A* was shown to inhibit sucrose export from the phloem to the apoplast through inhibition of SWEET-transporters (Abelenda et al., 2019) thereby enhancing the efficiency of sucrose delivery to sink tissues (van den Herik et al., 2021). Using a biophysical model of sugar and water transport we recently demonstrated that this dual role of *StSP6A*, tuber induction and inhibition of SWEET-mediated export, preferentially enhances sucrose allocation to the tuber sink (van den Herik & ten Tusscher, 2022). A remaining open question is what processes make tubers strong sinks, and to what extent *StSP6A* is involved in controlling these processes.

While *StSP6A* mediated induction of tuberization is a logical first step in establishing a new strong sink, whether and how the switch from apoplastic to symplastic unloading (Viola et al., 2001) contributes to sink-strength has remained unclear. Although symplastic unloading is generally believed to enhance unloading efficiency and thereby contribute to sink strength (Fernie et al., 2020; Viola et al., 2001), this is largely based on data comparing different species and tissues rather than the comparing the two distinct unloading modes for a single tissue. Additionally, the processes guiding this unloading switch and the potential role of *StSP6A* therein have not yet been elucidated. In addition to inhibition of the apoplastic route through *StSP6A* inhibition of SWEETs, a coordinated promotion of the symplastic route is likely to occur.

Plasmodesmal aperture, a key factor determining the efficiency of symplastic transport, is regulated by callose deposition at the neck of the plasmodesmata, with reduced callose deposition opening plasmodesmata. Callose homeostasis is regulated by two antagonistic gene families, callose synthases (CalS), producing callose and β-1,3-glucanase (1,3-BG) degrading callose (Amsbury et al., 2018; De Storme & Geelen, 2014; Wu et al., 2018). We therefore investigate whether upon tuberization onset changes in callose levels and the expression of callose homeostasis genes occur.

Another transition occurring during tuber formation is the switch from energy to storage metabolism. This switch involves a transition from apoplastic cell-wall invertase (cwInv) to cytoplasmic sucrose-synthase (SuSy) mediated sucrose cleavage (Viola et al., 2001). While the former delivers glucose and fructose for energy metabolism, the latter serves as an initial step in the formation of starch for storage by directly yielding UDP-Glucose, the precursor for starch synthesis (Nazarian-Firouzabadi & Visser, 2017). Potato tubers have been shown to depend on a switch to SuSy usage during tuber growth to prevent inefficient growth and storage (Bologa et al., 2003). During the early stages of tuber growth, a substantial increase in the hexose/sucrose ratio, and a decrease in total sugars occurs. The decrease in total sugars is indicative of an increased starch synthesis rate (Oparka et al., 1990), whereas a decreased hexose/sucrose ratio is likely due to increased rates of metabolism of hexoses derived from sucrose, coupled with an increased flux of sucrose into the developing tip (Davies, 1984; Ross & Davies, 1992). Altogether there is thus enzymatic and metabolic evidence for a functional enzymatic switch to occur. Still, an open question is whether this enzymatic switch is triggered by the change in sucrose delivery due to the apoplastic to symplastic switch, or rather is part of a more coordinated developmental program changing both transport mode and metabolism. We therefore investigate the extent of concurrence of the changes in callose and sucrose metabolism.

The switch to formation and storage of non-soluble starch and concurrent decrease in soluble sugars may enhance sink strength under symplastic unloading by maintaining a concentration gradient towards the sink. Alternatively, the increased sink-strength following the unloading switch may involve the differences between the transport modes themselves. Passive symplastic transport through plasmodesmata is generally regarded one or a few orders of magnitude more efficient than active apoplastic transport (Patrick & Offler, 1996). Moreover, symplastic transport reduces the energy spent on sucrose delivery by eliminating the energy required to maintain the proton motive force and reduces the requirement for significant investment in vascular tissue (Patrick & Offler, 1996). The relative efficiency of the two modes of transport will thus likely depend on the physiological conditions present in the stolon and tuber, and enhanced sink strength may involve both transport mode and metabolism. To investigate the coupling between transport mode and metabolism, we used a biophysical model to compare apoplastic versus symplastic unloading efficiency as a function of transporter and plasmodesmata densities, local sucrose concentration and concentration gradients.

## Materials & Methods

### *In vitro* plant growth and microscopy

Solanum Andigena was propagated *in vitro* on MS20 medium (MS, 20g/L saccharose, pH5.8), and cultivated at 24 °C for four weeks. The nodes from the upper three nodal stem sections were selected with each node containing an expanded leaf and a single axillary bud. At least 40 decapitated single-node cuttings were cut and propagated in MS20 medium under dark conditions at 20 °C for 5 to 10 days. Explants with elongated stolons were moved to tuber induction medium (MS, 80g/L saccharose, 1.5mL/L 6-Benzylaminopurin (BAP), pH5.8) and cultivated in similar conditions as stolon elongation.

For microscopy five samples were harvested at three different development stages; non-swelling stolon (stage 1), swelling stolon/small tuber (stage 2 or 3), large tuber (stage 4) as described by Viola et al. (2001). Harvested stolon and tuber samples were cut using a hand microtome. Samples were then stained in a 150mM K_2_PO_4_ (PH=9) and 0.01% aniline blue solution for 2 hours in the dark. Callose deposition was imaged with a DAPI filter (4FL) and an excitation wavelength of 370nm.

### Sequence retrieval, phylogenetic analysis and functional annotation

Protein sequences of the β-1,3-glucanase, Callose Synthase and Invertase families were identified using the Phytozome database (Goodstein et al., 2012) and a PSI-BLAST (Altschul et al., 1997) search was performed for each family to identify similar sequences missing or wrongly annotated in the Phytozome database. For each family, multiple sequence alignments were made with the auto option in MAFFT v7.310 (Katoh & Standley, 2013) and trimmed with trimAl v1.rev15 (Capella-Gutiérrez et al., 2009) using a gap threshold of 10%. Phylogenetic trees were constructed with IQ-TREE v1.5.5 (Nguyen et al., 2015) using ModelFinder (Kalyaanamoorthy et al., 2017) and ultrafast bootstrap approximation (Hoang et al., 2018).

The 1,3BG family was functionally annotated for features previously associated with the protein family, similarly to the approach used by Paniagua et al. (2021); Signal peptide (SP), glycosylphosphatidylinositol-anchors (GPI-anchors), and X8 (CBM43) domains were identified. Preduction of these features was done using SMART (Schultz et al., 2000) and Interpro (Mitchell et al., 2019). Signal peptides (SP) were further predicted using SignalP v5.0 (Almagro Armenteros et al., 2019), and presence of GPI anchor was further predicted using PredGPI (Pierleoni et al., 2008). Sequence signatures were considered present when predicted by two or more databases.

### *In silico* gene expression analysis

Microarray expression data of 24 tissue and organ samples from *S. tuberosum* Group Phureja DM 1-3 516 R44 (DM) was obtained from spudDB (Pham et al., 2020). These samples were a selection of all samples without applied stress and/or treatment free. Similarly, 15 samples from S. *tuberosum* Group Tuberosum RH89-039-16 (RH) (Zhou et al., 2020) were analyzed to confirm the observations in DM. The DM dataset contained two stolon and three tuber samples which were investigated in more detail. Analysis of the expression data was performed in python using the clustermap function from the seaborn package (Waskom, 2021), expression was normalized using Z-score normalization and hierarchical clustering was performed using the ‘ward’ method implemented in seaborn.

### Unloading model

Unloading rates for three different unloading modes are compared for varying sucrose concentration in the phloem (C_phloem_) and parenchyma (C_parenchyma_). Below we describe the models and parameter derivation, all parameter values are given in Table 1. Apoplastic or active unloading (I_a_) is described by a Michaelis-Menten term dependent on the phloem sucrose concentration (eq.1).

**Table 1.** Variables and parameters for the potato sucrose unloading model. Values in brackets represent the reported range of the parameter in potato.

| Parameter/Variable | Symbol | Value (range) | Units | Reference |
| --- | --- | --- | --- | --- |
| Sucrose concentration phloem | $C_{\text{Phloem}}$ | Variable | $\text{mol m}^{-3}$ | - |
| Sucrose concentration parenchyma | $C_{\text{Parenchyma}}$ | Variable | $\text{mol m}^{-3}$ | - |
| Maximum transport rate | $v_{\text{max}}$ | 0.93<br>(0.5-2) | $10^{-19} \text{mol } \mu\text{m}^{-2} \text{s}^{-1}$ | Chen et al., 2012 |
| Michaelis-Menten constant | $K_M$ | 50<br>(10-75) | $\text{mol m}^{-3}$ | Chen et al., 2012 |
| Plasmodesmata density | N | 4<br>(3-6) | $\mu\text{m}^{-1}$ | Oparka (1985) |
| Diffusion constant sucrose | D | 5e-10 | $\text{m}^2 \text{s}^{-1}$ | - |
| Cell wall thickness | l | 1<br>(0.3-1.2) | $\mu\text{m}$ | Gibson, (2012)<br>Reeve et al. (1973) |
| Plasmodesmata outer radius | $r_{\text{outer}}$ | 10 | nm | Ross-Elliot et al. 2017 |
| Desmotubule radius | $r_{\text{dt}}$ | 8 | nm | Ross-Elliot et al. 2017 |
| Cytoplasmic sleeve width | w | 2 | nm | Ross-Elliot et al. 2017 |
| Gas constant | R | 8.31 | $\text{Pa m}^3 \text{K}^{-1} \text{mol}^{-1}$ | - |
| Temperature | T | 293 | K | - |
| Partial molal volume sucrose | $V_{\text{suc}}$ | 0.22 | $\text{dm}^3 \text{mol}^{-1}$ | - |
| Dynamic viscosity of water | $\eta_0$ | 1.002 | $\text{mPa s}$ | - |

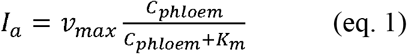

For this mode it is assumed that phloem unloading from the sieve-element/companion-cell (se-cc) complex is limiting, i.e. SWEET-transporter kinetics are the bottleneck in this step, we therefore parameterized this step with SWEET-specific parameters. Expected differences between SWEET and SUT-transporters facilitating uptake in the parenchyma are small (van den Herik et al., 2021). SWEET rates were reported as 39 ± 6 pmol/oocyte/min (Chen et al., 2012), which was rewritten to 0.9e-19 mol/μm^2^/s by using reported oocyte dimensions (Wallace & Selman, 1981). Reported SWEET K_m_ values range from 10-75mM (Chen et al., 2012), here an intermediate value of 50mM was used.

To simulate symplastic transport we used the model for simple plasmodesmata developed by Ross-Elliott et al. (2017). This model described two symplastic unloading modes; diffusive symplastic transport and symplastic bulk flow, both through plasmodesmata. Both models therefore need plasmodesmal area as important input. Individual plasmodesmal area is calculated by subtracting the surface area of the desmotubule from the total area, assuming that both the plasmodesmal and desmotubule surface area is circular. Total plasmodesmal area (A_PD_) is then calculated by multiplying individual area with the total number of plasmodesmata per μm^2^:

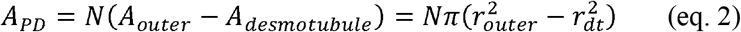

Diffusive unloading through plasmodesmata (I_D_) is described as standard diffusion through a simple collection of pores (eq. 3) with area A_PD_ and pore length l. Diffusion thus relies on the sugar concentration gradient between the phloem and parenchyma (Δc):

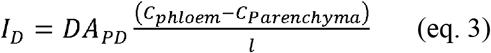

Bulk flow through plasmodesmata (I_b_) is described by the volumetric water flow through the plasmodesmata (Q) as well as the sugar concentration in the phloem (eq. 4).

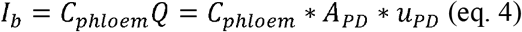

The volumetric water flow through plasmodesmata is driven by the pressure differential between the phloem and parenchyma, we here assume that this pressure differential is solely dependent on the osmotic pressure differential, and as such that turgor pressure in the phloem and surrounding parenchyma is equal (eq. 5)

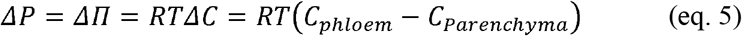

With R being the universal gas constant and T the absolute temperature.

Water flow velocity through plasmodesmata is estimated as flow through a straight slit of width *w (*where w = r_outer_ - r_dt_) and length l as described in detail by Ross-Elliot et al. (2017), resulting in equation 6.

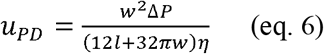

In this model version the viscosity of the solution flowing through the plasmodesmata (η) was solute dependent (eq. 7). Plasmodesmal sugar concentration was estimated as the average between phloem and parenchyma.

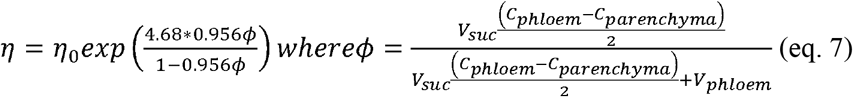

Thus, bulk transport is linearly dependent on phloem sucrose concentration (Eq. 4), linearly dependent on the phloem parenchyma sucrose concentration difference determining the osmotic pressure differential (Eq. 5), and non-linearly dependent on this same concentration difference via the viscosity of the phloem sap (Eq. 7).

## Results

### Callose deposition at the phloem decreased during tuberization

To investigate whether callose levels decrease during tuber formation we used fluorescence microscopy, using aniline blue to stain callose. Callose was visualized in stolon and tuber samples, with clear presence of callose in both phloem rings in the stolon (Fig. 1A, B). Xylem vessels were also clearly visible due to the fluorescent properties of lignin (Albinsson et al., 1999). Callose abundance was strongly decreased in the vascular region of tuber samples (Fig. 1C,D). Precise quantification of callose in tubers was prohibited by the large amount of starch interfering with the callose signal, preventing a more quantitative comparison between stolon and tuber callose levels. Nonetheless, a decrease in callose levels from the stolon to tuber stage was consistently observed in all samples (n=5 stolons and tubers).

**Figure 1.**
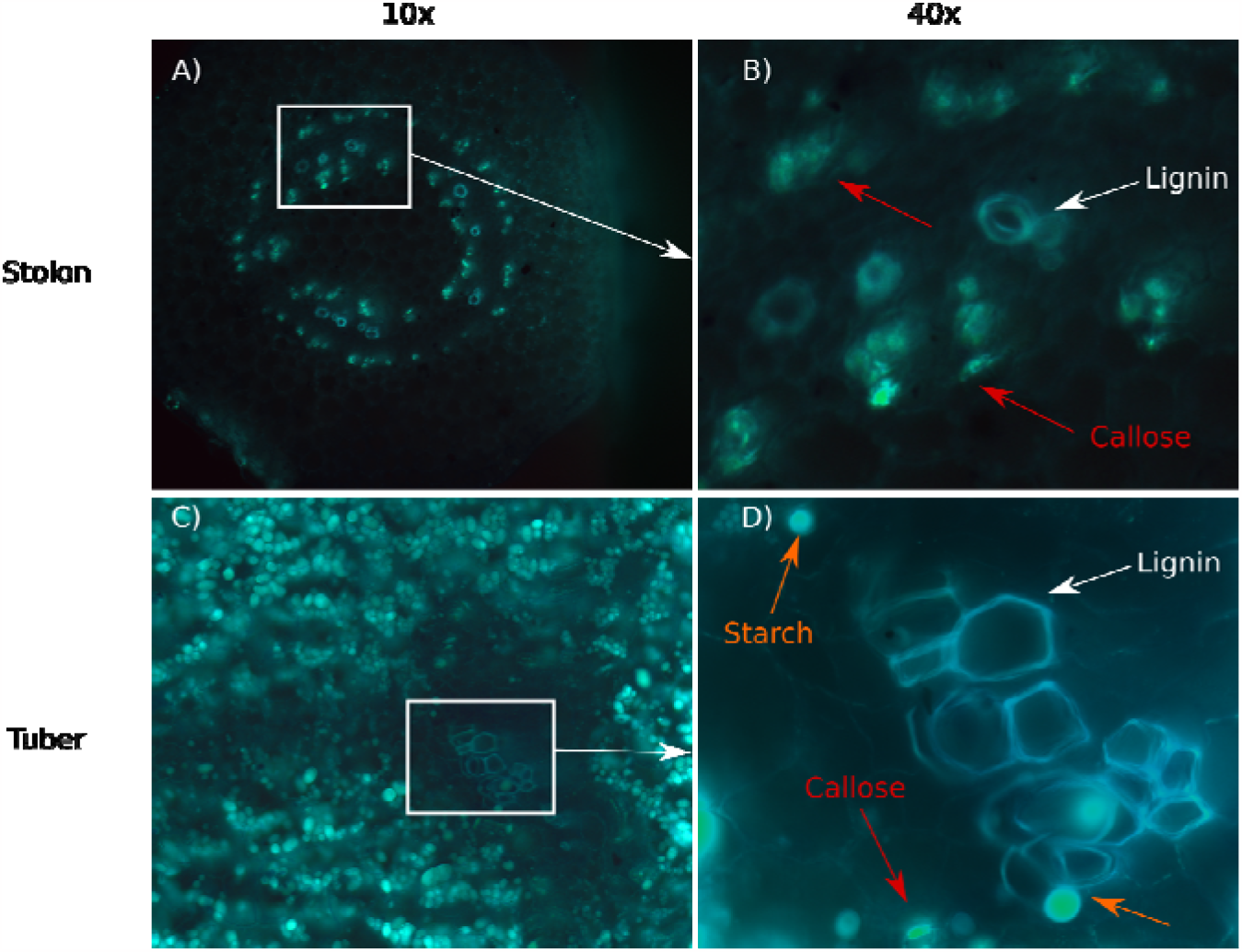
Callose presence at stolons and tubers. A) Longitudinal sample of a non-swelling stolon at 10x magnification B) 40x magnification of part of the sample in A. Callose (red arrows) and lignin (white arrows) are clearly visible in the phloem and xylem C) Longitudinal sample of the vasculature of a large tuber (stage) at 10x magnification D) 40x magnification of part of the sample in C. Callose (red arrows), lignin (white arrows) and starch (orange arrows) are visible in the phloem and xylem Sequence retrieval and phylogeny of the 1,3-BG, CalS and invertase gene families

### Sequence retrieval and phylogeny of the 1,3-BG, CalS and invertase gene families

To investigate gene expression patterns, a classification of sugar and callose metabolism gene families was needed. While the SuSy (Van Harsselaar et al., 2017; Xu et al., 2019), SWEET (Manck-Götzenberger & Requena, 2016) and SUT (Chen et al., 2022; Chincinska et al., 2008) families were previously classified (Table 2), an overview of potato 1,3-BG, CalS and sucrose invertase genes was lacking. We first identified the genes constituting these families and performed a phylogenetic and functional analysis to get an overview of the number of genes (Table 2) present and their expected functional role in tuber development.

**Table 2.**
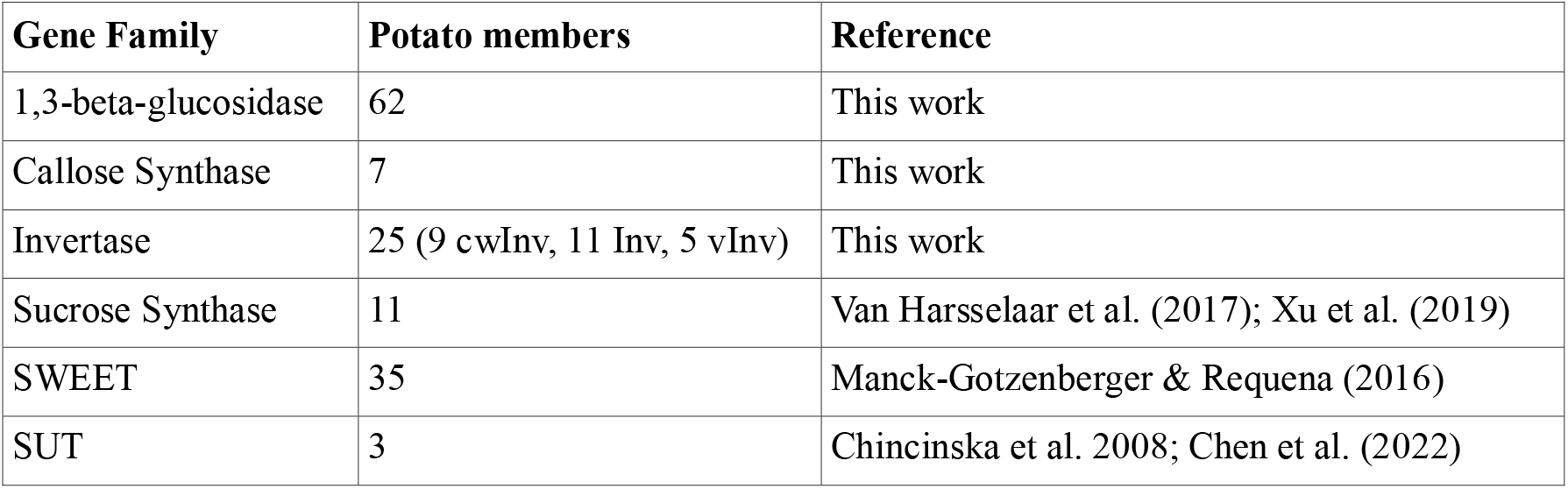
Overview of the callose and sucrose gene families.

| Gene Family | Potato members | Reference |
| --- | --- | --- |
| 1,3-beta-glucosidase | 62 | This work |
| Callose Synthase | 7 | This work |
| Invertase | 25 (9 cwInv, 11 Inv, 5 vInv) | This work |
| Sucrose Synthase | 11 | Van Harsselaar et al. (2017); Xu et al. (2019) |
| SWEET | 35 | Manck-Gotzenberger & Requena (2016) |
| SUT | 3 | Chincinska et al. 2008; Chen et al. (2022) |

A total of 62 1,3-BG genes encoding callose degrading enzymes were identified using the phytozome database combined with a psi-BLAST search. Together with a group of 8 Arabidopsis 1,3-BG genes previously identified to be plasmodesmata-associated (Levy et al., 2007; Wu et al., 2018) containing one or more representatives of each of the 3 previously identified clades (α,β,γ) (Doxey et al., 2007) a phylogenetic tree was constructed (Fig. S1). Genes were functionally annotated for the presence of a signal peptide (excretion), GPI-anchor (membrane anchorage) and a X8-domain (carbohydrate binding/plasmodesmata association). Our results indicate that the three previously described clades are all present in potato and differ significantly in the presence/absence of functional domains. Specifically, the α-clade is very diverse with regards to the presence of the domains, while the β-clade is characterized by presence of an excretion signal peptide, GPI-anchor, and X8-domain in most proteins (13 out of 17), and the γ-clade is characterised by a complete absence of GPI-anchor and X8-domain, similar to what is found in Arabidopsis (Doxey et al., 2007) and tomato (Paniagua et al., 2021). Based on this, proteins active in callose degradation at the PD are expected to be mainly localized in the α- or β-clade. Indeed, all 1,3-BG characterised to be PD-related in *Arabidopsis* are located in the α-clade (Levy et al., 2007).

A total of 13 CalS genes encoding callose producing enzymes were identified, 6 sequences were discarded as they lacked the UDP-glucose catalytic site needed for enzymatic activity (Hong et al., 2001). The 7 remaining CalS sequences, together with the 13 CalS genes reported in Arabidopsis (Richmond & Somerville, 2000) were used to construct a phylogenetic tree (Fig. S2). The tree was annotated with known functions for the Arabidopsis genes (Wu et al., 2018), inferring similar roles for potato genes from their *Arabidopsis* orthologs. Based on this analysis two putative PD-associated proteins were identified (Soltu.DM.07G023050 & Soltu.DM.01G001920).

Finally, with regards to invertases involved in sucrose degradation, three main clades exist, acidic cell wall invertase (cwInv), neutral/alkylic soluble invertase (Inv), and vacuolar invertases (vInv). These three classes have largely different physiological roles (Roitsch & González, 2004). A total of 25 invertase genes were identified in potato. Not all genes were functionally annotated, and function was therefore inferred using their phylogeny and partial annotation in potato (Fig. S3), based on this analysis 9 cwInv, 11 Inv and 5 vInv are present.

### The highly diverse 1,3-BG and CalS families show clear developmental expression clusters

We next investigated the expression patterns of the 1,3-BG and CalS families to investigate changes in callose homeostasis during tuberization. For this we made use of a dataset containing microarray gene expression data across a variety of organs of *S. tuberosum* Group Phureja DM1-3 (DM) (Pham et al, 2020). Distinct developmental expression clusters are present, with flowers and stamens, petioles, fruit, stolons and tubers all clustering separately (Fig. S4). Stolons did show partially similar expression patterns to shoot and root samples, which is expected from the shared stem-like nature of these three organ types. The observed clusters reflect the functional diversity of callose besides its role in regulation of the plasmodesmal aperture, as callose also plays a role in cell wall integrity and mechanics, response to (a)biotic stresses, pollen development and cellular differentiation (Levy et al., 2007; Wu et al., 2018; Amsbury et al., 2018).

Subsequently, three tuber and two stolon tissue samples available in the DM-dataset were investigated in more detail, In these five samples a subset of 52 of the 62 1,3-BG genes and all 7 identified CalS genes were expressed (Fig. 2). Stolon samples clustered together, with all CalS genes active in stolon samples and none showing high activity in the tubers. Tuber samples were also strongly correlated, with two showing very strong overlap in 1,3-BG expression and the third sample having a separate set of 1,3-BGs expressed. Tuber samples mainly expressed 1,3-BGs from the γ-clade, lacking both a GPI-anchor as well as a X8-domain, with only 2 out of the 21 1,3 BG genes having a GPI-anchor (Fig. 2). In contrast, in stolons expression of α-, and β-clade is dominant and 17 out of 31 1,3-BG genes contained a GPI-anchor domain. The majority of 1,3-BGs active in stolons are thus membrane and/or PD-associated, suggesting a role in callose degradation. In contrast in tubers, expressed genes are associated with pathogen resistance and cell-wall remodeling (Doxey et al. 2007), likely due to their secretory nature. To verify these observations, the same analysis was performed on a stolon and tuber expression dataset of *S. tuberosum* Group Tuberosum RH89-039-16 (RH) (Zhou et al., 2020), providing similar results (Fig. S5). Combined, this shows that callose synthase is mainly active in stolons, whereas 1,3-BG is active in both stolons and tubers, albeit that functionally different genes are expressed.

**Figure 2.**
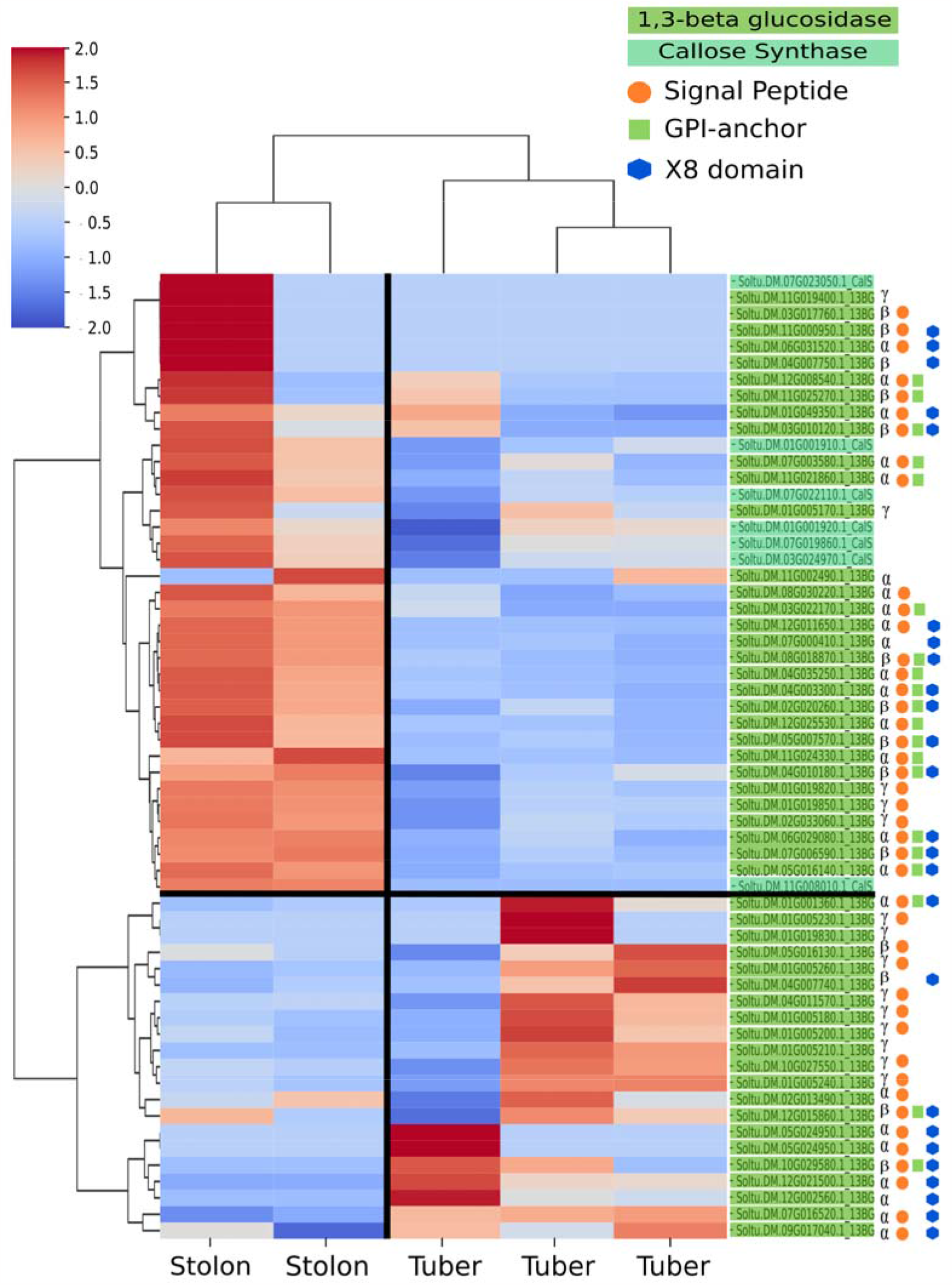
1,3-BG and CalS expression in stolon and tuber samples. Z-score normalized rows with hierarchical clustering shows distinct stolon and tuber clusters. Clade and annotation of signal peptides, GPI-anchors and X8-domains is visualized behind each 1,3-BG gene.

### A clear developmental pattern is present for callose and sucrose metabolism

Besides changes in callose homeostasis, a rewiring of sucrose metabolism as well as changes in sugar transporters are known to occur during tuberization (Viola et al., 2001; Jing et al., 2023). To investigate if and how these processes are coupled during tuberization the gene expression of callose (1,3-BG & CalS) and sucrose metabolism (SuSy, cwInc, vInv & Inv) as well as sucrose transporters (SWEET & SUT) were investigated alone, as well as in combination (Fig. 3, S6). Clearly distinct stolon and tuber clusters were observed when considering sucrose metabolism in isolation (Fig. S6), as well as combined with sucrose transporters and callose metabolism (Fig. 3).

**Figure 3.**
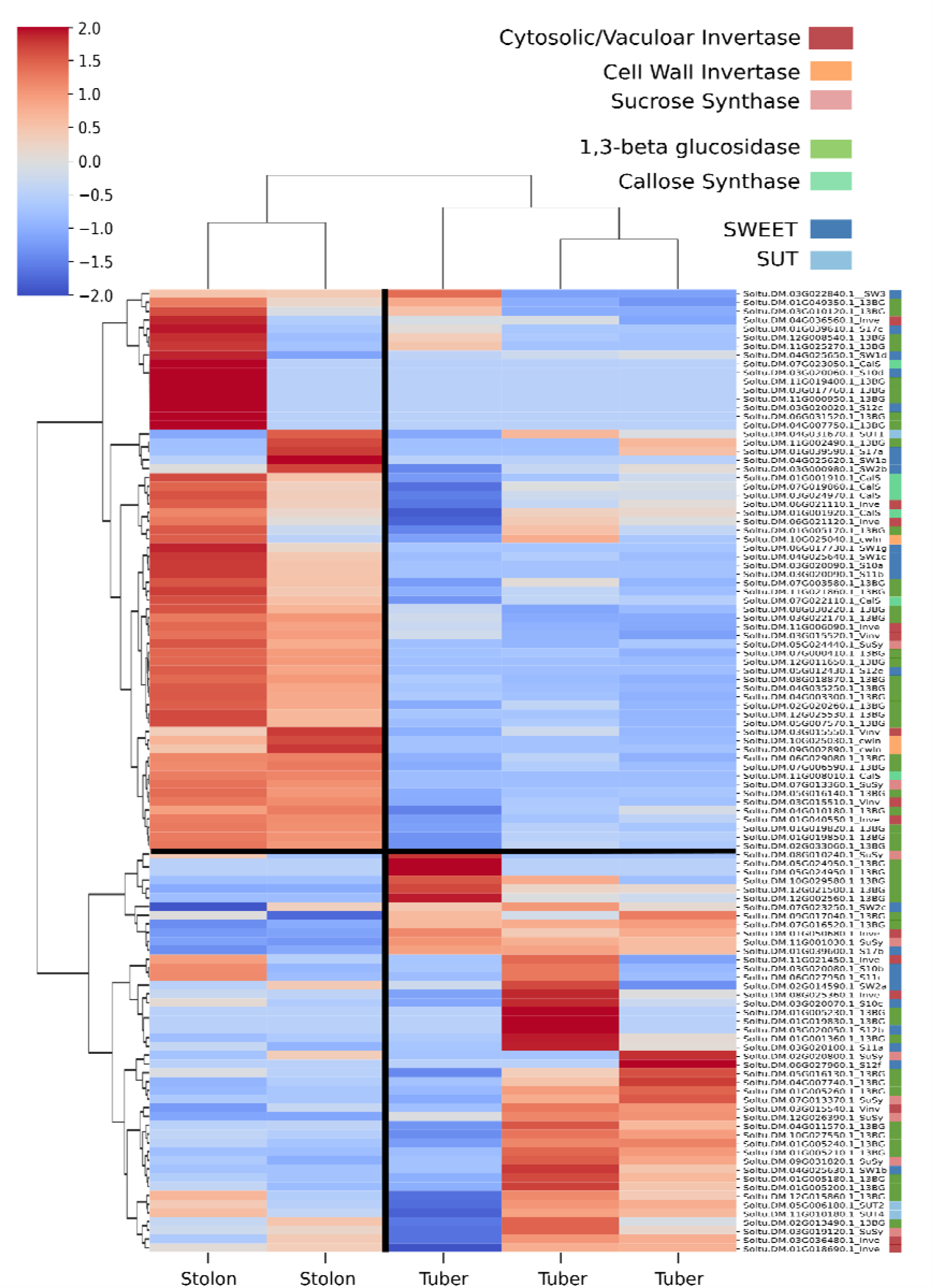
Heatmap of expression profiles of sugar metabolism, transport and callose balancing in stolon and tuber samples of DM. Z-score normalized rows with hierarchical clustering show similar stolon and tuber clusters as observed in Fig. 2. Colors behind the genes depict gene family.

During the developmental switch from stolon to tuber both cytoplasmic and cell wall invertase activity is reported to decrease, whereas SuSy activity increases (Viola et al., 2001). Three out of nine cwInv are expressed in the stolon cluster and show no to very low expression levels in tubers (Fig 4, S6). The other six cwInv were not expressed in either stolons or tubers. Soluble and vacuolar invertases showed more mixed expression patterns, with five active in the stolon cluster, and six active in the tuber cluster. Nine SuSy genes were expressed, with the majority active in tubers (seven out of nine). SWEET and SUT expression patterns were more ambiguous than sucrose and callose metabolism, consistent with earlier observations (Jing et al., 2023), and a more important role for post-translational interactions such as between *StSP6A* and *StSWEET11b* is expected for these families.

**Figure 4.**
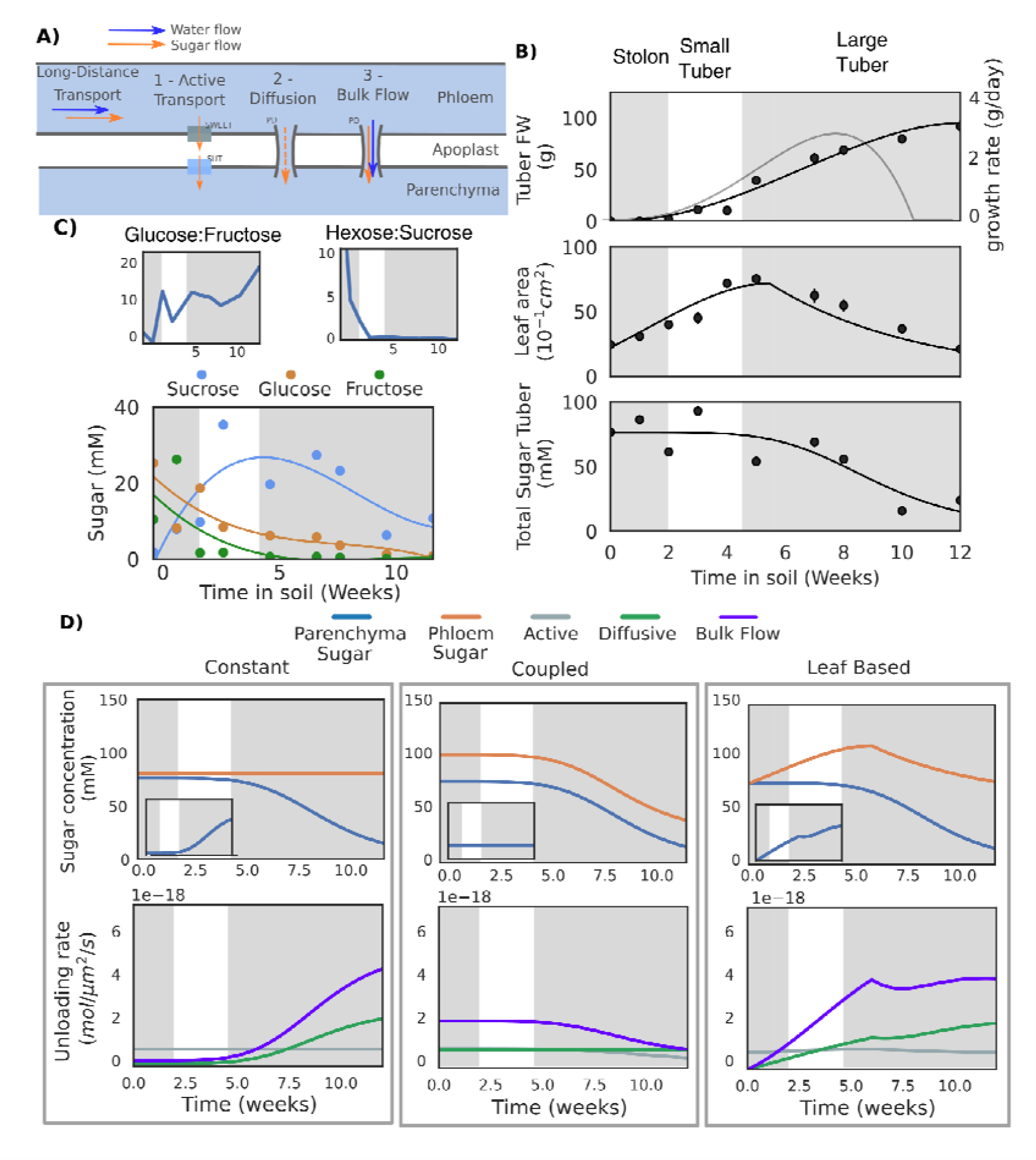
Sucrose unloading under different physiological conditions in stolon and tuber. **A)** Schematic overview of the unloading model. **B)** Potato tuber fresh weight, leaf area and tuber sugar dynamics over plant development (van den Herik et al. 2023). The first grey box depicts the stolon stage, white small tuber stage and the second grey large tuber stage, as described by Viola et al. (2001) **C)** Sucrose, fructose and glucose dynamics in stolons and tuber (van den Herik et al., 2023). Glucose:Fructose and Hexose:Sucrose ratios are calculated from the same data. **D)** Model results for the three phloem concentration scenarios (columns). The top row shows the sucrose levels in the phloem and parenchyma (see B), with the inset showing the gradient between the phloem and sucrose. The bottom row gives the unloading rates for the three unloading modes as obtained from the simulations.

### Enzymatic changes create favored conditions for passive unloading

Above we demonstrated how the switch from stolon to tuber involves concurrent changes in both sucrose and callose metabolism, both of which may contribute to the formation of a strong tuber sink. Here we investigate the relative importance of sucrose physiology versus unloading mode in determining sink strength. We first describe the physiological changes in terms of phloem and parenchyma sugar concentrations. Combined with estimated sucrose transporter levels and plasmodesmata density, we parameterized a simple apoplastic and symplastic unloading model developed by Ross-Elliot et al. (2017) for the stolon and tuber (Fig. 4A) (See methods).

Due to the low K_m_ of SWEET-transporters (50mM) apoplastic transport is close to saturation under most conditions, except for the coupled scenario, where lower concentrations during tuber bulking cause transport to decrease too half the maximum rate during development (Fig 4D, grey lines). SUC transporters have an even lower K_m_ of 10mM, and operate close to saturation for all tested concentrations. As a consequence, apoplastic transport is largely constant across concentrations and scenarios. In contrast, diffusive and bulk flow unloading do show large differences over development and between the scenarios. As diffusion is only dependent on the concentration gradient between the phloem and parenchyma the unloading rate parallels the concentration gradient (Fig 4D, insets and green lines). Plasmodesmal bulk flow depends on both the sugar concentration gradient which drives osmotic pressure and hence flow rate, and the concentration of sucrose transported in the phloem. As a consequence, bulk flow increases more strongly with concentration gradient, resulting in a higher unloading potential than diffusion under all conditions, as was also observed in the developing phloem of Arabidopsis (Ross-Elliott et al., 2017).

Our model demonstrates that the switch to storage metabolism that results in a decrease in tuber sugar levels potentiates diffusive and bulk symplastic unloading in both the conservative, constant and data-based, leaf-surface driven scenarios where sugar concentration gradients increase. In both cases, bulk flow significantly exceeds diffusive transport due to the non-linear dependence of bulk flow on concentration levels via osmotic pressure and convective solute flow (Fig. S7) In contrast, in the coupled scenario, where concentration gradients remain constant while phloem sucrose levels decrease, diffusive unloading efficiency remains constant while bulk unloading efficiency declines over development. This scenario is however deemed unlikely given the observed leaf growth dynamics, as well as the known negative feedback regulation of sucrose levels on photosynthesis which likely prevents a fully parallel decline of phloem sucrose levels with sink levels.

Importantly, since we estimated transporter expression levels, plasmodesmal densities and apertures, model outcomes depend on precise parameter settings. To test the robustness of our results we performed simulations with 2-fold higher transporter levels, 2-fold lower plasmodesmal densities and these changes combined (Fig. S8). Our results show that while the precise rates of transport and the cross-over point from whereon symplastic transport is more efficient shifts, no qualitative changes occur. Put differently, we consistently observe that during early tuber formation apoplastic transport is more efficient, while during later stages symplastic transport dominates. The model thus clearly demonstrates that the physiological conditions created by the enzymatic switch (decreasing parenchyma concentration due to increasing starch synthesis) and general plant dynamics (leaf area increase) unlocks the higher potential of passive unloading.

## Discussion

In this study, we explored the significance of the apoplastic to symplastic unloading switch and the transition from energy to storage sugar metabolism on tuber sink-strength increase during and after tuberization. We first investigated whether, in addition to the previously identified StSP6A mediated decrease in apoplastic transport upon tuberization, also plasmodesmata further opened. For this we used callose deposition as a proxy for plasmodesmata opening. Using fluorescence microscopy, we showed that callose levels did decrease in tuber samples, indicating an important role for decreased callose deposition in the unloading switch.

With changes in callose deposition confirmed, we continued with investigation of gene expression changes of callose homeostasis genes, i.e. Callose Synthase for synthesis and β-1,3-glucanase for degradation. We observed that while 1,3-BGs from the γ-clade were expressed in tuber samples, expression of α-, and β-clade is dominant in stolons. In *Arabidopsis* the γ-clade proteins have been associated with pathogen resistance and cell-wall remodeling (Doxey et al. 2007). Increased activity of the excreted γ-clade proteins might thus be an indicator of faster cell-growth and has been associated with fast growth in pollen tubes (Wang et al., 2022). The majority of 1,3-BGs active in stolons are membrane and/or PD-associated, suggesting a role in callose degradation. Expression patterns in stolons thus suggest high degradation potential at plasmodesmata, while in tubers this potential is decreased and growth-, and pathogen-associated expression dominates. Phylogenic inference of function for the smaller CalS family revealed clear expression of gametophyte associated proteins in the flower/stamen cluster (Soltu.DM.11G008010) and two putative PD-associated proteins (Soltu.DM.07G023050 & Soltu.DM.01G001920), which were highly expressed in fruit, callus, and stolon samples. No to low expression of CalS in tubers was present. Combined, this analysis suggests that callose homeostasis at plasmodesmata is a constant process of synthesis and degradation in stolons, as also observed in *Arabidopsis thaliana* pollen tubes, where callose is transiently present (Abercrombie et al., 2011). While in tubers the transient presence of callose at plasmodesmata is replaced by low callose deposition due to decreased synthesis and relocation of degradation to the apoplastic space. It thus indicates that decreased callose deposition in tubers is caused by decreased synthetic CalS activity and not increased 1,3-BG activity. Recently, Nicolas et al. (2022) showed that both downregulation of CalS and upregulation of 1,3-BG resulted in increased symplastic movement in aerial tubers. There, the downregulation of CalS was again more prominent than the upregulation of 1,3-BG activity.

We next investigated the concurrence of the changes in callose and sucrose metabolism. Expression of CalS and cwINV are exclusively clustered in stolon samples, whereas SuSy expression dominates in tubers. Low expression of SuSy genes was present in stolons, which can possibly be explained by the need for its product UDP-glucose in both stolons and tubers as it is a shared precursor for both starch and callose metabolism (Barnes & Anderson, 2018). Furthermore, low individual expression levels can be caused by technical or biological noise between samples in this dataset. Overall, this indicates a well-coordinated developmental switch, with a combined transition from growth (cwInv) to storage (SuSy) metabolism, and from callose PD homeostasis to extracellular callose degradation.

Lastly, we set out to understand the implications of this coordinated switch on the unloading potential and thus sink-strength of tubers. To this end, we parameterized a biophysics-based phloem unloading model (Ross-Elliot et al., 2017) for stolons and tubers. Using this model, we demonstrated that it is the combined switching of the unloading mode and the sucrose metabolism that increases tuber sink strength. The metabolic switch ensures maintenance of the concentration gradient necessary for efficient symplastic unloading, while starch metabolism is further activated because of increased cytoplasmic sucrose inflow (Stein & Granot, 2019; Winter & Huber, 2000). Clearly the finding that passive gradient driven transport increases with an enhanced gradient is in itself trivial. Additionally, the exact time point at which symplastic transport exceeds apoplastic transport in efficiency of course depends on the precise parameterization of apoplastic and symplastic transport rates and densities. The key point of our current modeling effort lies in the demonstration that -unless an unrealistic phloem concentration scenario is applied-during initial stolon and tuber development apoplastic transport is more efficient while at later stages symplastic transport is more optimal. This underlines that 1) the *in planta* observed switch in transport mode is physiologically sensible, 2) symplastic transport is not always more optimal. Indeed, in fruit plants sucrose storing fruits and seeds typically depend on a switch to apoplastic unloading to prevent a symplastic back-flow of the soluble sugars down the created concentration gradient (Ma et al., 2019). In the current model, enzyme levels were kept constant, as e.g. different SWEET subtypes showed different expression dynamics upon tuberization. Additionally, we kept enzyme activity constant, despite the previously demonstrated *StSP6A* mediated decline in *StSWEET11* activity.

Thus, in reality, active apoplastic transport would decrease in later stages in all scenarios, further favoring symplastic transport. On a similar note, we kept PD density and area constant, while we showed that callose removal led to an increased openness and thus diameter. Other processes, such as increasing PD-densisty over development or changes in other gating proteins, shown to increase PD conduction (Lucas et al., 2009) were also not considered. These changes would have further favored symplastic over apoplastic unloading. Finally, we here used ‘simple’ PD architecture, shown to be up to 10x less effective than funnel plasmodesmata (Ross-Elliot et al., 2017). Overall, the model used here is thus a conservative, worst-case, scenario. As such, the model strongly supports that it is the combined switching of sugar metabolism and unloading mode that enables an increased tuber sink-strength.

## Acknowledgements

We thank Sam von der Dunk and Julian Vosseberg for help with phylogenetic tree construction and interpretation. Furthermore, we thank Johan Buchner for help with microscopy.

## Author contributions

BH performed phylogenetic analysis, gene expression analysis, model construction and analysis of the models. SB designed the *in vitro* growth protocol and performed microscopy. YL grew the plant material, performed microscopy and did the first phylogenetic and gene expression analysis. CB conceived the project and analyzed the data. KT conceived the project, analyzed the data and performed the model construction and analysis. All authors approved the submitted version.

## Data availability

No data was generated in this study.

