## Supplemental_Figures for "A coordinated switch in sucrose and callose metabolism enables enhanced symplastic unloading in potato tubers"

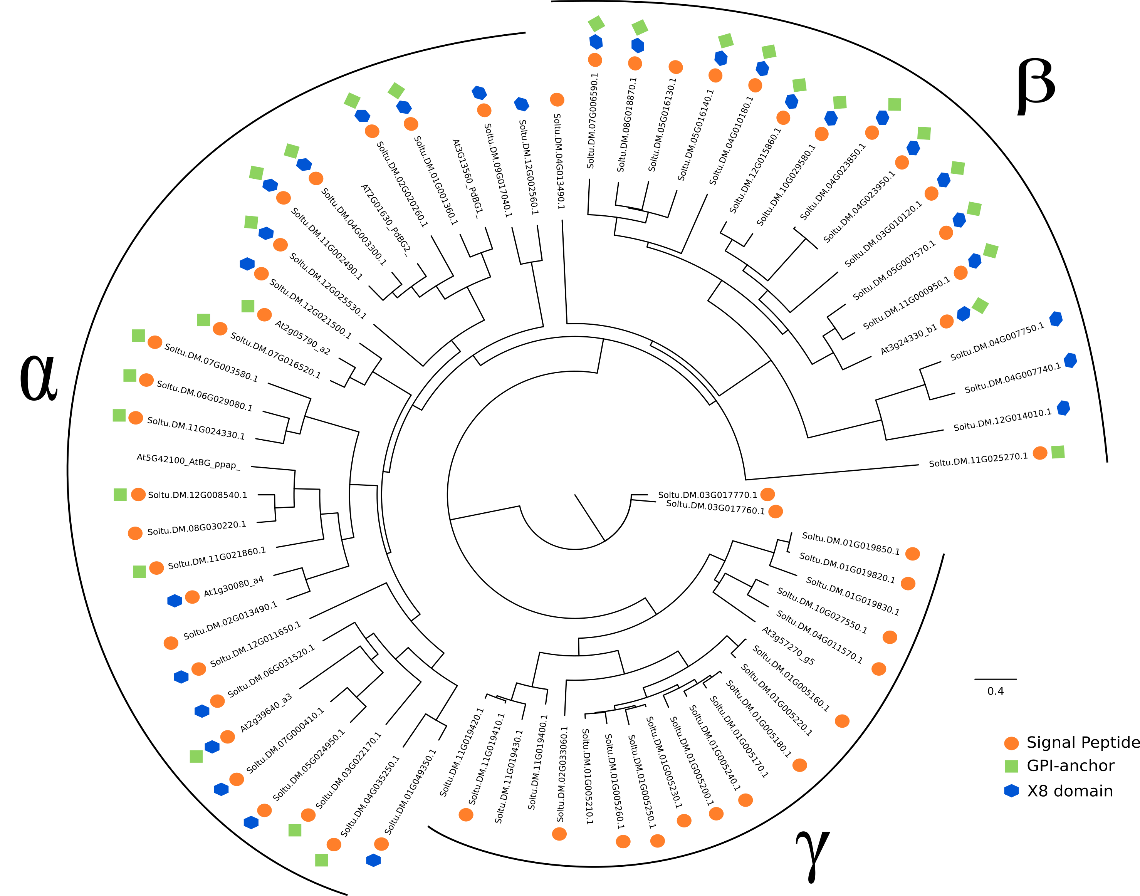

**Figure S1. Phylogentic tree of the Beta-1,3-glucosidase family annotated for plasmodesmata related domains.** Presence of three functional domains was depicted using symbols next to the gene names. The three clades were highlighted using α, β, and γ as previosuly described by Doxey et al. (2007).

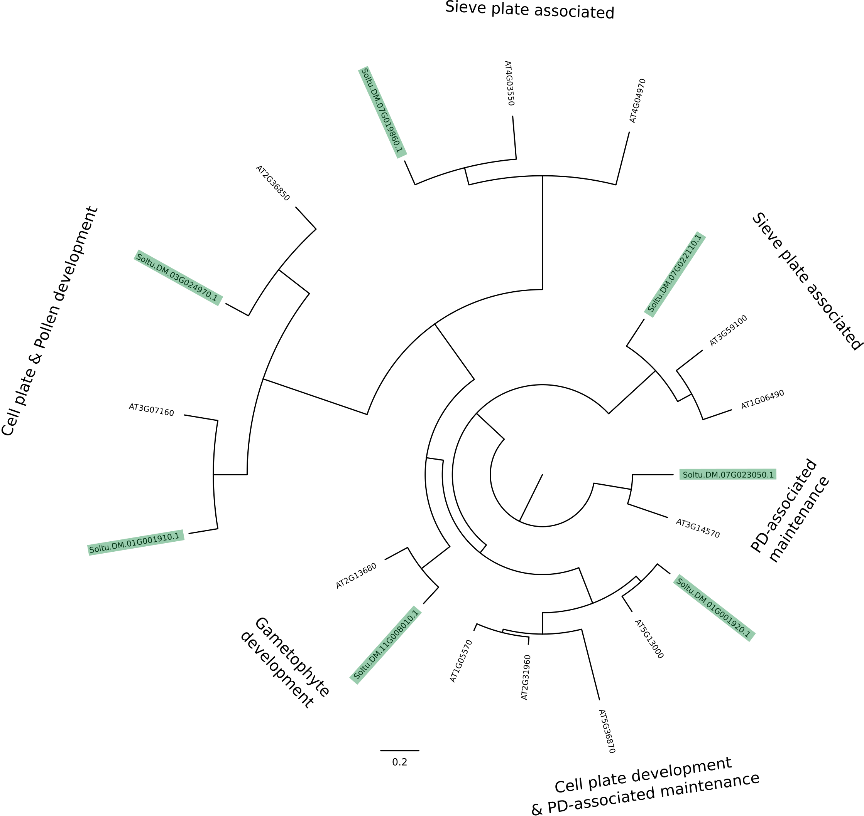

**Fig S2. Phylogentic tree of the Callose Synthase family functionally annotated using function derived from the A. thaliana orthologs.** Annotation of arabidopsis genes was obtained from Wu et al. (2018). Genes highlighted in green were present in the potato genome, non-highlighted genes were annotated genes from Arabidopsis.

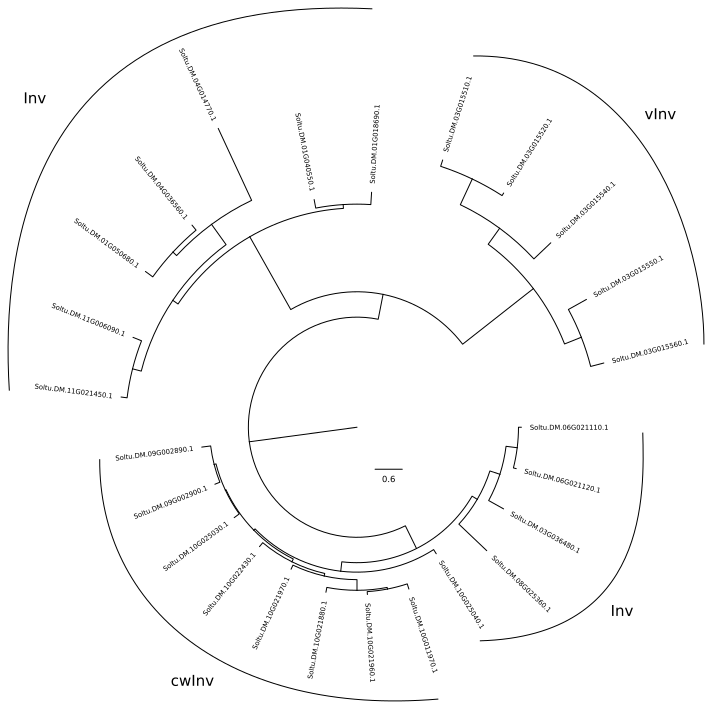

**Figure S3.** **Phylogenetic tree of the invertase gene family.** Not all invertase genes found via BLAST search were annotated. Therefore, phylogenetic placement in the tree above was used to functionally annotate the invertase genes that were not annotated as cell wall (cwInv), soluble (Inv) or vacuolar invertase (vInv).

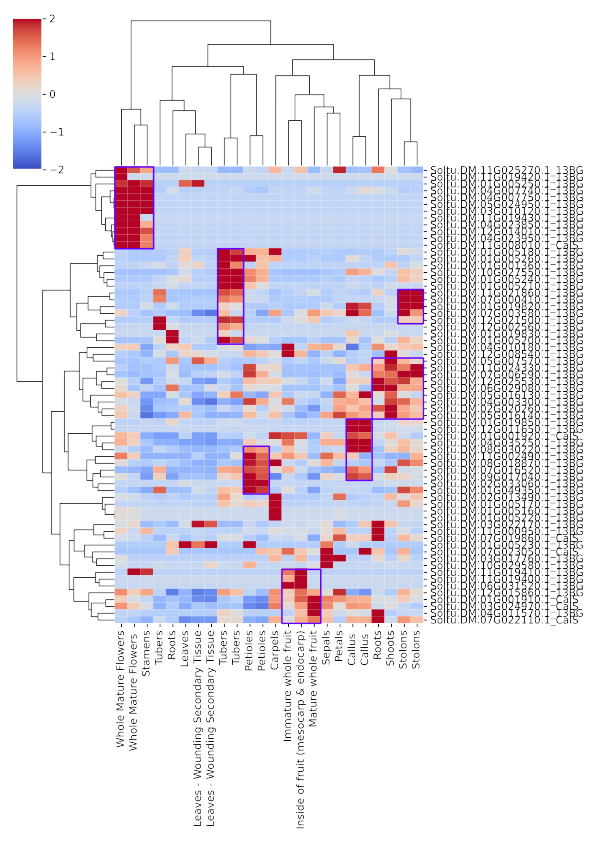

**Fig S4. Beta-1,3-glucosidase and Callose Synthase family expression in above- and belowground samples reveal clear developmental patterns.** Purple boxes show high expression clusters in specific samples.

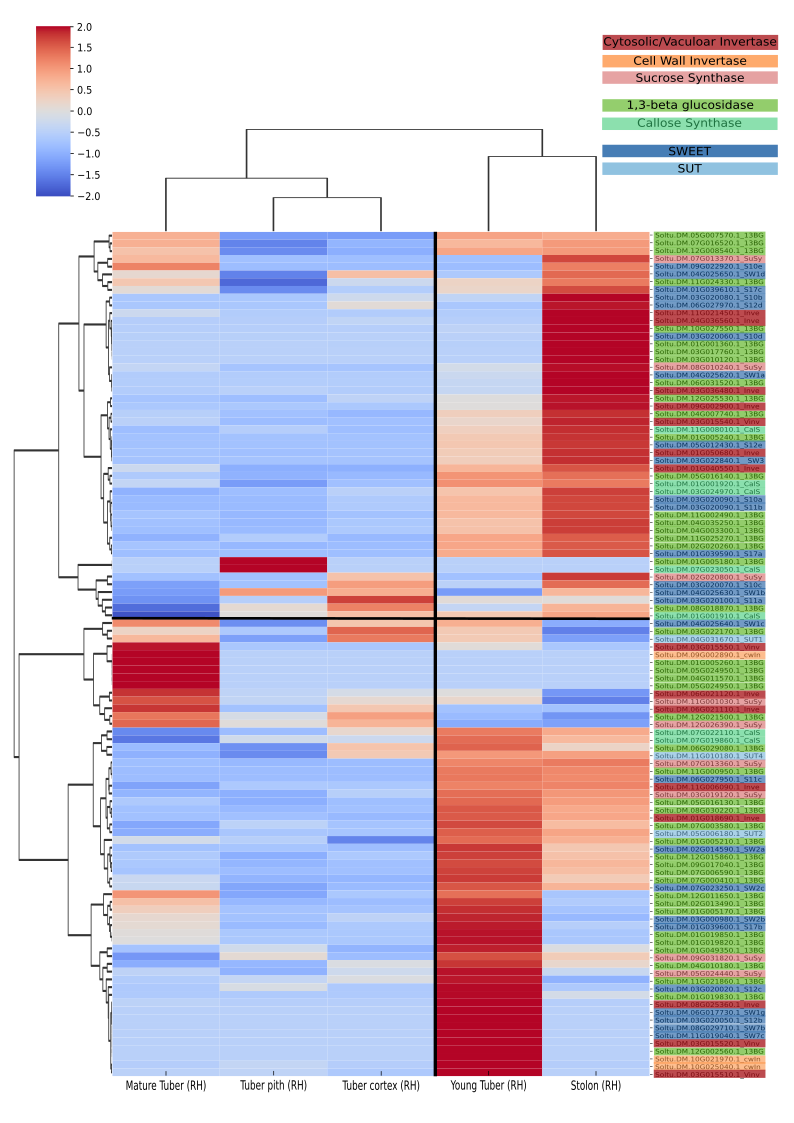

**Figure S5. Callose, sucrose and transporter gene families expression in tuber and stolon samples in *S. tuberosum* Group Tuberosum RH89-039-16.** Clear developmental cluster are again present for stolons and tubers. In the RH dataset, there is a distinction between young and mature tubers, which do show distinct expression patterns. Patterns are less clear than in DM. Overall, we do again see expression of CalS in stolons and young tubers, and cwInv expression lowest in mature tubers.

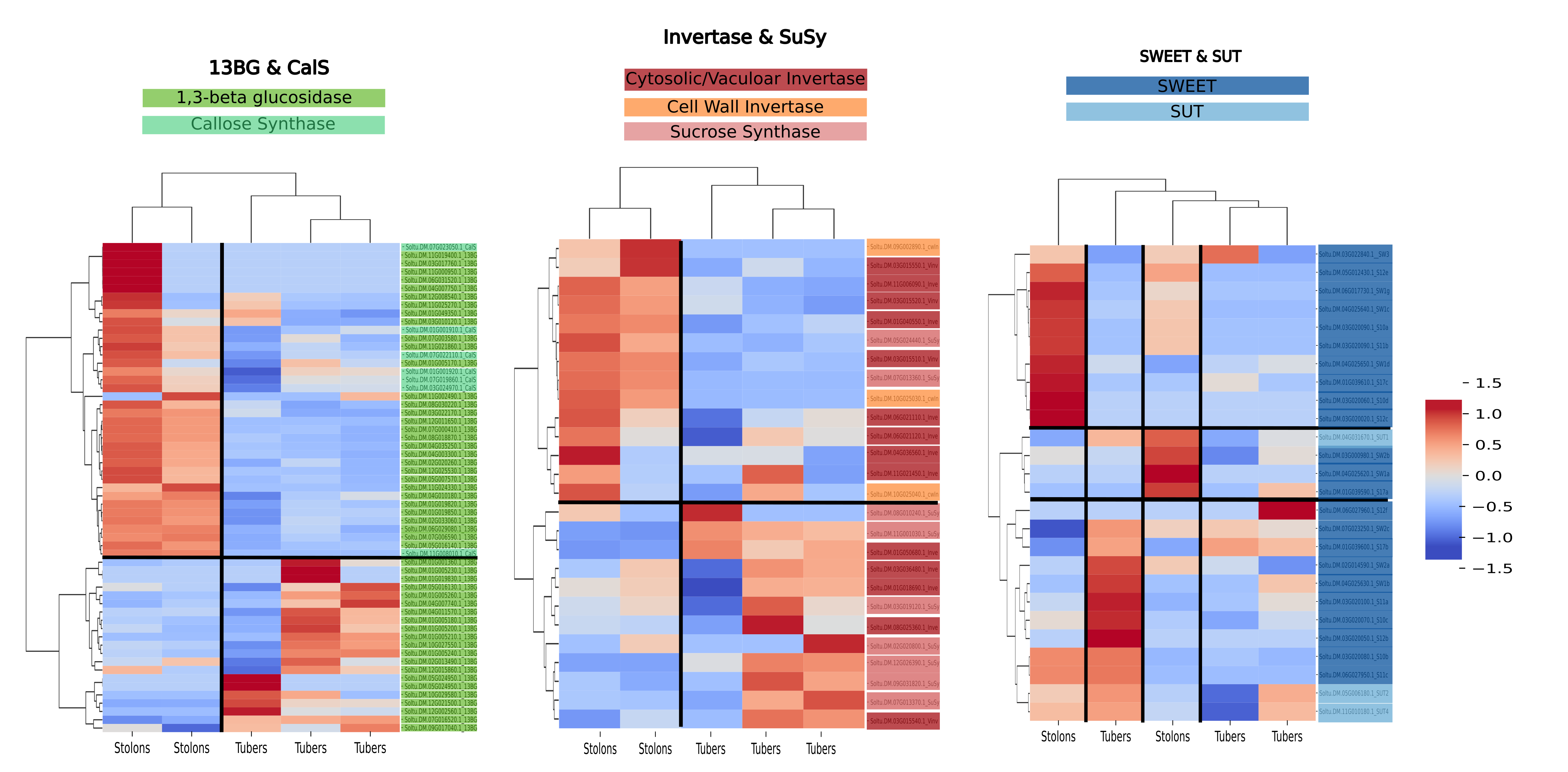

**Figure S6. Gene families clustered per functional category in DM samples**. These three heatmaps show clear developmental clusters for tubers and stolons in Sucrose and Callose metabolism. On the other hand, SWEET and SUC transporters show more ambiguous expression patterns.

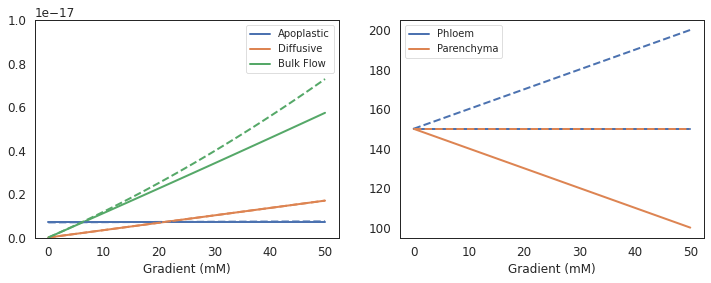

**Fig S7. Unloading rates as a function of the concentration gradient.** Dotted lines represent a gradient caused by an increasing phloem concentration and solid lines a gradient caused by decreasing parenchyma concentration.

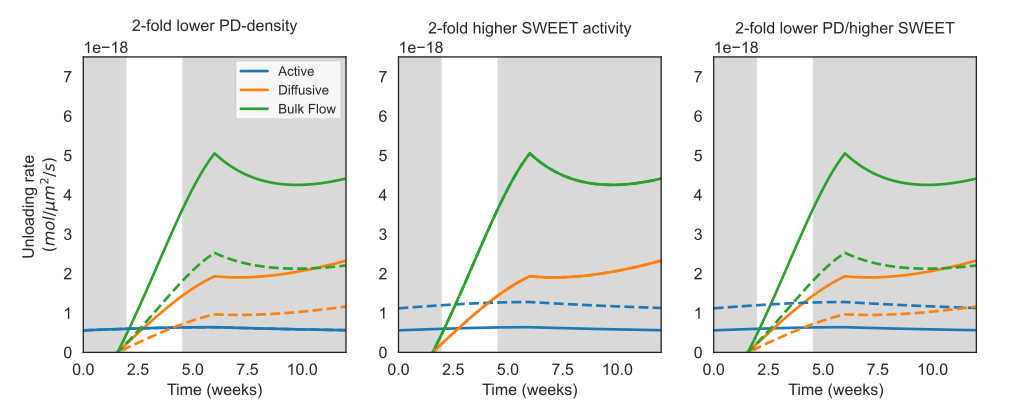

**Fig S8. Robustness of the unloading model to PD-density and SWEET-activity changes.** Phloem conditions for these simulations were leaf-based and all parameters were equal to those used in Fig. 4D, except PD-density and SWEET-activity, as shown in the figure titles. Solid lines are baseline results equal to results in Fig. 4D, dashed lines represent simulation results with altered parameters. These plots show that while the exact quantitative results change, qualitative behaviour of the model does not for realistic parameter regimes.
